# CRISPR/Cas9 gene editing in *Drosophila* via visual selection in a summer classroom

**DOI:** 10.1101/2024.03.28.587232

**Authors:** Lutz Kockel, Valentina Zhang, Jenna Wang, Clara Gulick, Madeleine E. Laws, Arjun Rajan, Nicole Lantz, Ayla Asgarova, Lillian Dai, Kristian Garcia, Charlene Kim, Michelle Li, Patricio Ordonez-Acosta, Dongshen Peng, Henry Shull, Lauren Tse, Yixang Wang, Wenxin Yu, Zee Zhou, Anne Rankin, Sangbin Park, Seung K. Kim

## Abstract

CRISPR/Cas9 methods are a powerful *in vivo* approach to edit the genome of *Drosophila melanogaster*. To convert existing *Drosophila GAL4* lines to *LexA* driver lines in a secondary school classroom setting, we applied the CRISPR-based genetic approach to a collection of *Gal4* ‘driver’ lines. The integration of the *yellow*^+^ coat color marker into homology-assisted CRISPR knock-in (HACK) enabled visual selection of *Gal4-to-LexA* conversions using brightfield stereo-microscopy available in a broader set of standard classrooms. Here, we report the successful conversion of eleven *Gal4* lines with expression in neuropeptide-expressing cells into corresponding, novel *LexA* drivers. The conversion was confirmed by *LexA-* and *Gal4-*specific GFP reporter gene expression. This curriculum was successfully implemented in a summer course running 16 hours/week for seven weeks. The modularity, flexibility, and compactness of this course should enable development of similar classes in secondary schools and undergraduate curricula, to provide opportunities for experience-based science instruction, and university-secondary school collaborations that simultaneously fulfill research needs in the community of science.

## Introduction

Controlled gene expression in *Drosophila melanogaster* is a powerful way to elucidate functions of cells, tissues, and genes in development and metabolism, and is principally accomplished through binary systems (Brand and Perrimon 1993; Lai and Lee 2006; Potter et al 2010; Kim et al 2021). These use a combination of a gene-specific promoter driving the expression of a *trans*-acting transcriptional activator (like *GAL4, LexA, or QF*) that regulates target transgene(s) expression through a *cis*-acting DNA element specific for that transactivator. This approach can be extended by combining two or more independent binary expression systems to query inter-organ or inter-cell interactions. (Lai and Lee 2006; Bosch et al. 2015, Gordon and Scott 2009; Bosch et al. 2015, Macpherson et al. 2015, Tsao et al 2022; reviewed in Kim et al 2021).

The generation of multiple binary expression systems has been limited by the relative paucity of specific driver lines. While several thousand stocks of *Gal4* lines are available, the number of publicly-available *LexA* driver lines remains comparatively low, though efforts have begun to address this resource gap (Pfeiffer et al 2010, Kockel et al 2016; Kockel et al 2019; Kim et al 2023). Recently, homology-assisted CRISPR knock-in (HACK) and similar CRISPR/Cas9-based methods were built to convert the large number of existing *Gal4* lines into driver lines expressing *QF* or *LexA* (Lin and Potter, 2016, Chang et al 2022; Karuparti et al 2023). In this method, HACKing inserts the *T2A*.*LexA* DNA into the *Gal4* ORF. This disrupts the *Gal4* gene, and results in T2A-mediated expression of *LexA* from the very locus of the HACKed *Gal4*.

In prior work, the identification of successfully ‘HACKed’ *Gal4*-to-*LexA* convertants required fluorescence microscopy (Lin and Potter, 2016). A modified and successful approach to convert existing *Gal4* lines into *LexA* drivers was recently described as a convenient tool-kit suitable for an 11-week secondary school class (Rankin et al., 2023); in that work, integration of a *yellow*^+^ cassette as a selectable marker gene permitted brightfield microscopy-based methods to identify successful *Gal4*-to-*LexA* convertants. We postulated that this technical innovation could broaden use of this experimental approach (Rankin et al., 2023).

Here, we applied a modular “CRISPR curriculum” to a summertime seven-week long course. This was an extension of similar efforts by Stanford University investigators to integrate experimental *Drosophila* genetics (hereafter, the Stan-X curricula) into high schools and colleges (Kockel et al., 2016). We report the successful implementation of the CRISPR curriculum into a summer school setting in 2023. Students of the class successfully converted eleven neuropeptide-*Gal4* lines into corresponding *LexA* drivers. The conversion was confirmed by *LexA*- and *Gal4*-specific GFP reporter gene expression.

## Methods

### *Drosophila* strains

The following fly stocks were obtained from the Bloomington *Drosophila* Stock Center (BDSC). *Gal4* lines: *w*; Burs-Gal4* (40972), *w*^*1118*^; *Capa-Gal4* (51969), *w*^*1118*^; *Proc-Gal4* (51972), *w*^*1118*^; *AstA-Gal4* (51979), *w*^*1118*^; *ETH-Gal4* (51982), *w*^*1118*^; *Mip-Gal4* (51984), *w*^*1118*^; *Lk-Gal4* (51993); *w*^*1118*^; *FMRFa-Gal4* (56837), *w*; NPF-Gal4* (25681), *y,w*; Akh-Gal4* (25683), *y,w*; CCAP-Gal4* (25685), *w*; Aug21-Gal4* (30137).

“Tool” stocks, including *w*; 10xUAS-CD8::GFP* in attP2 (32185), *w*; 13xLexAop-CD8::GFP* in attP2 (32203), *y*^*1*^,*w*^*67c23*^,*hsp70-Cre; CyO / Sco* (766), *y*,^*1*^,*w*^*1118*^ (6598). *y*^*1*^,*w*^*\**^, *vas-Cas9; CyO*^*HACKy*.*V2,y+,RFP+*^*/L*, and *y,w; CyO/L; TM2/TM6B* were previously described (Rankin et al., 2023).

### Conversion of *GAL4* to *LexA*^*G4H*^ by Cas9, and floxed *RFP*^*+*^,*y*^*+*^ cassette deletion from converted *LexA*^*G4H,y+,RFP+*^ lines

The intercrossing scheme is outlined in **Figure 1**. For F_0_, two males of the *GAL4* line and six virgin females of *y*^*1*^,*w*, vas-Cas9; CyO*^*HACKy*.*V2,y+,RFP+*^/ *L*, the *LexA* donor line, were crossed. Forty independent single *w*^+^ and *CyO F*_1_ male progeny was mated to two virgin *y*,^1^,*w*^1118^ females. *F*_2_ male progeny were screened for *y*^+^,*w*^+^ and *CyO*^-^ conversion events, and their RFP expression was confirmed. From these males, stocks were established (**Figure 1**). Single promoter-*LexA*^G4H^ conversion males were mated to *y*^1^,*w*^67c23^,*hsp70-Cre; CyO / Sco* without heat shock. A single *w*^+^ male offspring was then mated to *y,w; CyO/L; TM2/TM6B* virgin females. Afterwards, a single male *w*^+^,*y*^-^,*RFP*^-^ offspring was mated to a balancer line to establish the balanced stock (**Figure 1**).

**Figure 1:**
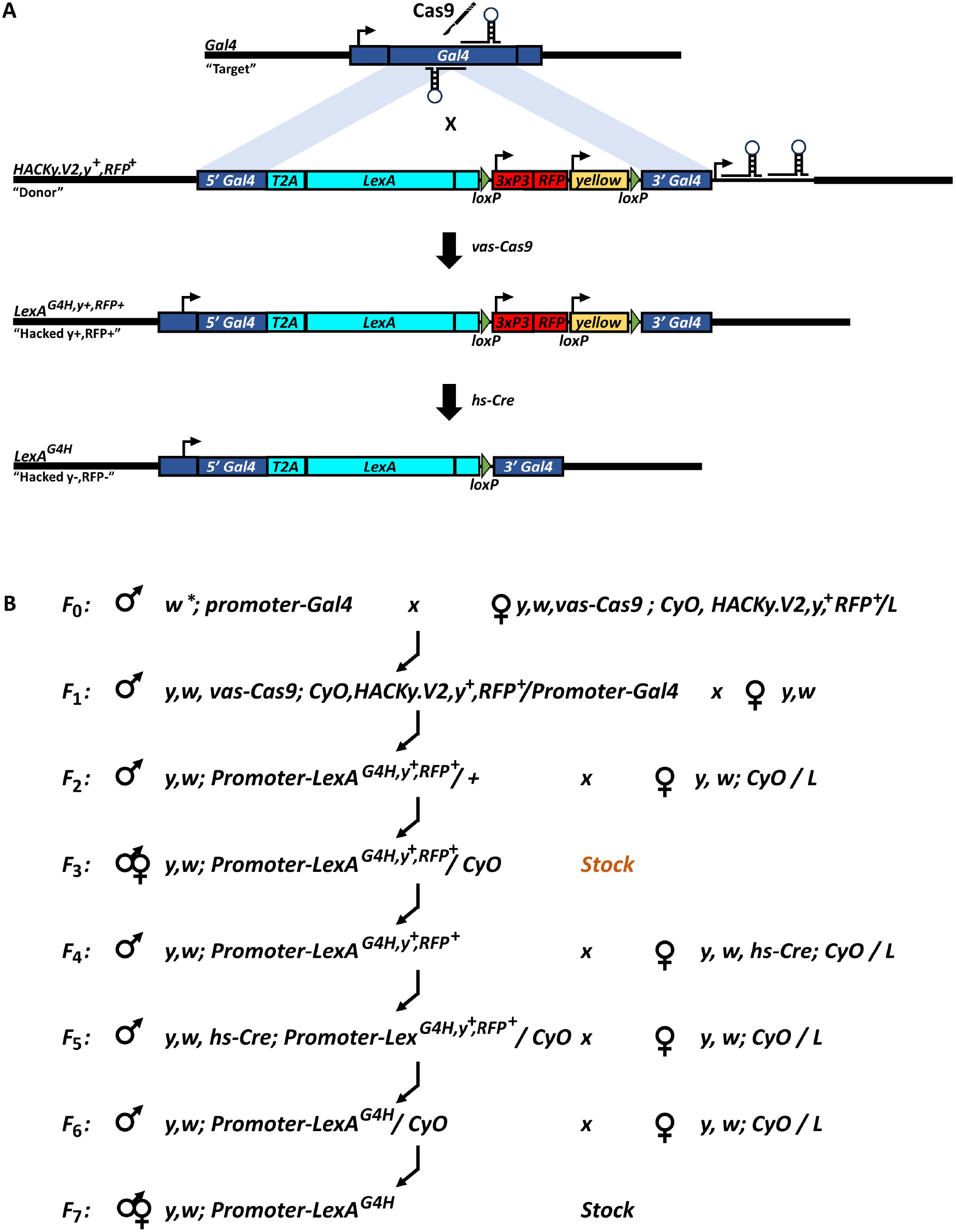
Genetic flow chart outlining HACKing (Homology Assisted CRISPR Knock-in) of *Gal4*. **(A)** Top: Crossing the target *Gal4* (dark blue, top construct) to the Version 2 (V2) donor strain encoding Cas9 controlled by the vasa-promoter on X (not shown), and the donor construct on *CyO* containing T2A.*LexA* marked by the floxed 3xP3-RFP, yellow+ cassette, flanked by *Gal4* homology arms and the U6-driven guide RNAs (*CyOHACKy*.*V2,y*^*+*^,*RFP*^*+*^). 3rd line from top: The resulting HACKed chromosome where the *Gal4* ORF has been disrupted and replaced by *T2A*.*LexA*, marked by the visual markers *yellow+* and *RFP+*. Bottom: The *yellow*^*+*^,*RFP*^*+*^ cassette is removed after a cross to *hs-Cre*. **(B)** Genetic crossing scheme to conduct *Gal4* hacking using the *CyO*^*Hacky*.*V2,y+,RFP+*^ donor, followed by removal of the floxed *y*^*+*^,*RFP*^*+*^ cassette by *hs-Cre*. Selection of individual HACKed males marked by *y*^*+*^,*RFP*^*+*^ takes place in *F2*. The HACKed stock is established in *F3*, the goal of the course. *RFP*^*+*^,*y*^*+*^ cassette removal to *F7* was conducted after the course had ended.

### Imaging *UAS-GFP* and *LexAop-GFP* reporter gene expression

Females of *w*; 10xUAS-CD8::GFP* and *w*; 13xLexAop-CD8::GFP* were mated to original target *Gal4* and converted *LexA*^*G4H*^ pairs to compare the resulting GFP reporter expression patterns. Inverted wandering third instar larvae were fixed in 4% paraformaldehyde in PBS for 30 mins, and fixed again in 4% paraformaldehyde in PBS containing 0.2% Triton X-100 overnight and rinsed and washed three times each in 1x PBS. L3 larval brains and L3 midguts were dissected from the washed carcasses, and mounted (SlowFade Gold antifade reagent with DAPI, invitrogen S36939) using Zeiss-made-by-Schott No. 1.5 170+/-5 um, 18 x 18 mm coverslips (Zeiss 0109030091). Images of GFP (recorded in green) and DAPI (recorded in blue) channels were recorded on a Zeiss AxioImager compound Epifluorescence microscope, captured using AxioImager software, and edited using Adobe Photoshop software.

### Course and Classroom organization

The *Drosophila Gal4* genome editing class took place in summer 2023 as a seven week ‘Animal Transgenesis” course in the Harvard University Department of Continuing Education (DCE). The class was held at the Biolabs of Harvard University, with instruction by personnel from Stanford University School of Medicine, The Lawrenceville School (NJ), and by two teaching assistants from Phillips Exeter Academy and the Lawrenceville School, both previously educated through the Stan-X program (Kim et al., 2023). The classroom was staffed with one-to-two instructors and two TAs every day. 12 participants, predominantly high school students, were selected by instructors.

The workflow of the course is detailed below in Results. In short, the first two weeks introduced *Drosophila* genetics and molecular biology of CRISPR/Cas9 genome editing, while later weeks were spent almost exclusively with hands-on work. The introduction was taught using a course-specific teaching manual covering the specific syllabus of the class. Class participation, written homework, midterm and final exams, and a final oral presentation by each student to their peers and instructors were an integral part of the curriculum and grading.

To accommodate the compression of experiments of the seven-week class, the *F*_0_ and *F*_1_ crosses were prepared ahead of the class start and handed to students for analysis after they had set up analogous crosses of *F*_0_ and *F*_1_. This compression of the ‘generation time’ for offspring enabled incorporation of the ‘*Gal4* / *LexA* expression module’ to study reporter gene expression within the course timeframe.

### Data and reagent availability

All converted *Gal4* derivatives are available at the Bloomington Drosophila Stock Center. Course manual is available on request.

## Results

### Selecting successful *Gal4*-to-*LexA* gene conversions *in vivo* using *yellow*^+^

We postulated that the HACK (Homology Assisted CRISPR Knock-In; Lin and Potter, 2016) crossing scheme using the visible selectable marker *yellow*^+^ (Rankin et al., 2023) could be adapted for use in an accelerated summer classroom setting lasting seven weeks (**Figure 1A, B**). Twelve neuropeptide-*Gal4* lines from the Bloomington *Drosophila* Stock Center (BDSC; Methods) were selected to convert to *LexA* driver lines in the 2023 “Animal Transgenesis” course at Harvard Summer School with 12 participants. The ‘version 2’ HACKing donor for *inter*-chromosomal conversion, HACKy.V2 located on the CyO chromosome (**Figure 1**; “CyO^HACKy.V2,y+RFP+^” donor hereafter), was reported to show optimized results for HACKing of second chromosome locations (Rankin et al., 2023), and the selected *Gal4* lines were reported to reside on chromosome II by their associated data in the BDSC database (see below).

Class participants successfully HACKed 11 of 12 *Gal4* lines. The exception, *AstA-G4* (Methods), was found to reside on chromosome III and was therefore not amendable to HACK-based conversion by the chromosome II CyO^HACKy.V2,y+RFP+^donor construct. The *yellow*^+^ selectable marker was sufficient to identify single males harboring the *Cas9*-mediated, *HACKy*.*V2,y+RFP+* dependent *Gal4*-to-*LexA* conversion event in the *F1* offspring (**Figure 1B**). Additional testing for the co-expression of the *RFP*^+^ eye marker in *yellow*^+^ conversion males showed both selectable markers were consistently present (**Figure 2**). Eye expression of RFP was driven by a *pax6* promoter element (Rankin et al., 2023; Sheng et al., 1997). In summary, within seven weeks, students successfully applied CRISPR targeting, genetics, and marker selection (Rankin et al., 2023) to derive multiple novel *LexA* lines.

**Figure 2:**
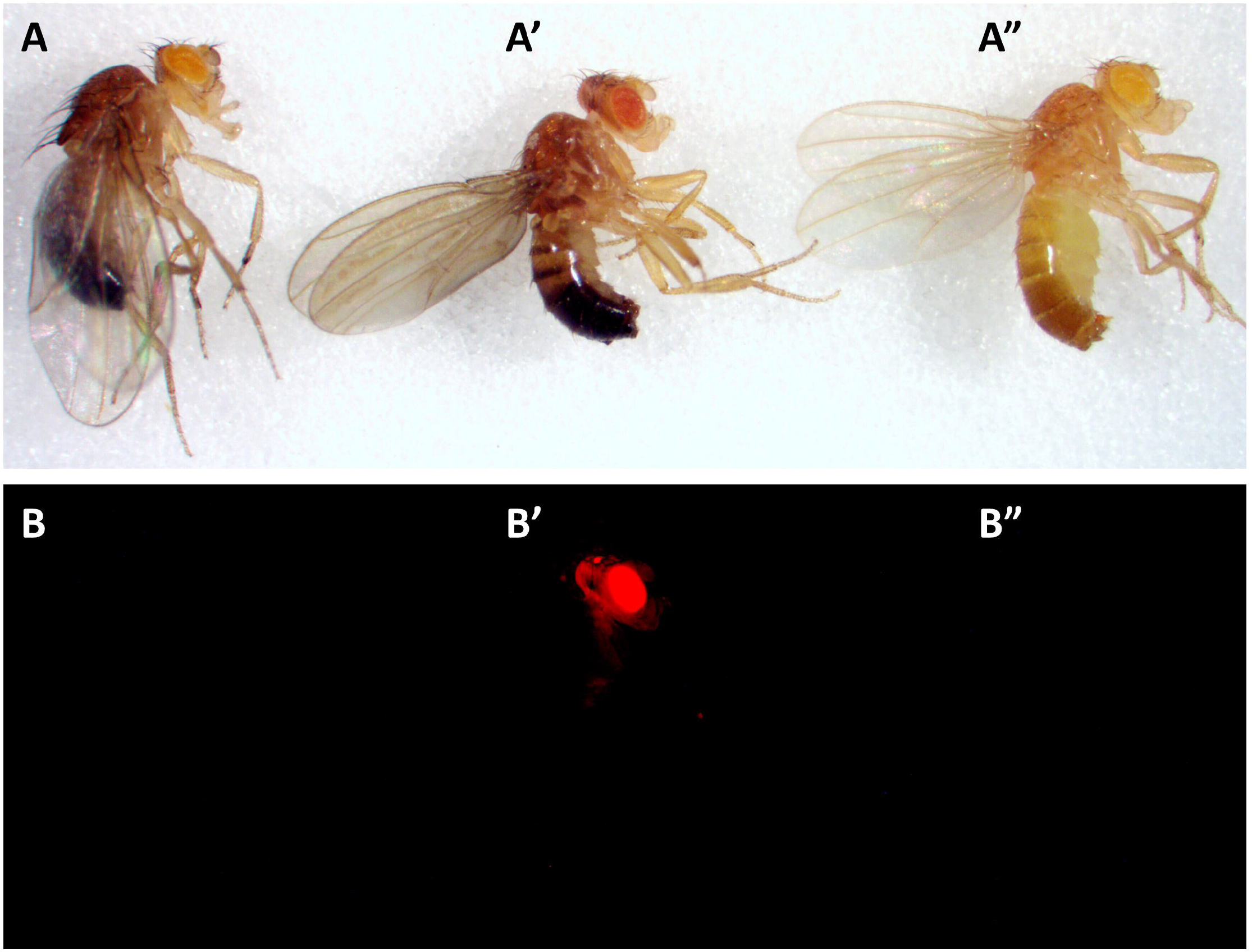
Expression of the *yellow+ and RFP+* reporters by the *LexA*^*G4H,y+,RFP+*^ intermediate only **(A-A”)** Stages of *Capa-Gal4* conversion analyzed via *yellow*^*+*^ reporter expression by brightfield microscopy. (A) *w*^*1118*^, *Capa-Gal4*, (A’) *y*^*1*^,*w*^*1118*^; *Capa-LexA*^*G4H,y+,RFP*^, (A”) *y*^*1*^,*w*^*1118*^; *Capa-LexA*^*G4H*^. Note that the parental *Capa-G4* line is in a white background only. **(B-B”)** Stages of *Capa-Gal4* conversion analysis of 3xP3-RFP^+^ reporter expression by epifluorescence microscopy. (B) *w*^*1118*^, *Capa-Gal4*, (B’) *y*^*1*^,*w*^*1118*^; *Capa-LexA*^*G4H,y+,RFP*^, (B”) *y*^*1*^,*w*^*1118*^; *Capa-LexA*^*G4H*^. Red: RFP.

### Comparison between parental *GAL4* and HACKy-derived *LexA* line expression patterns

The *in vivo* insertion of the *T2A*.*LexA* coding DNA into the *Gal4* ORF predicts the functional equivalence between the parental *Gal4* and the derivative *LexA*^*G4H,y,+RFP+*^ and the *LexA*^*G4H*^ expression domains. To test preservation of the expression domains, we crossed (1) the parental Gal4 to reporter flies harboring *10xUAS-mCD8::GFP*, and (2) converted *LexA*^*G4H,y,+RFP+*^ and *LexA*^*G4H*^ flies to reporter flies harboring the *13xLexAop2-mCD8::GFP* reporter transgene, then visualized GFP expression in L3-stage larval brains (**Figure 3**). Both GFP reporter genes are inserted into *attP2* and showed no transcriptional effect due to insertion position (Pfeiffer et al 2010, Rankin et al., 2023). To further examine the influence of the *yellow*^+^, *RFP*^+^ reporter cassette on the expression domain of *LexA*, we performed parallel intercrosses to investigate the expression of *LexA* where the *yellow*^+^, *RFP*^+^ cassette was either present (*LexA*^*G4H,y+,RFP+*^ **Figure 3A’**)or removed by hs-Cre (*LexA*^*G4H*^, Figure 3A”).

**Figure 3:**
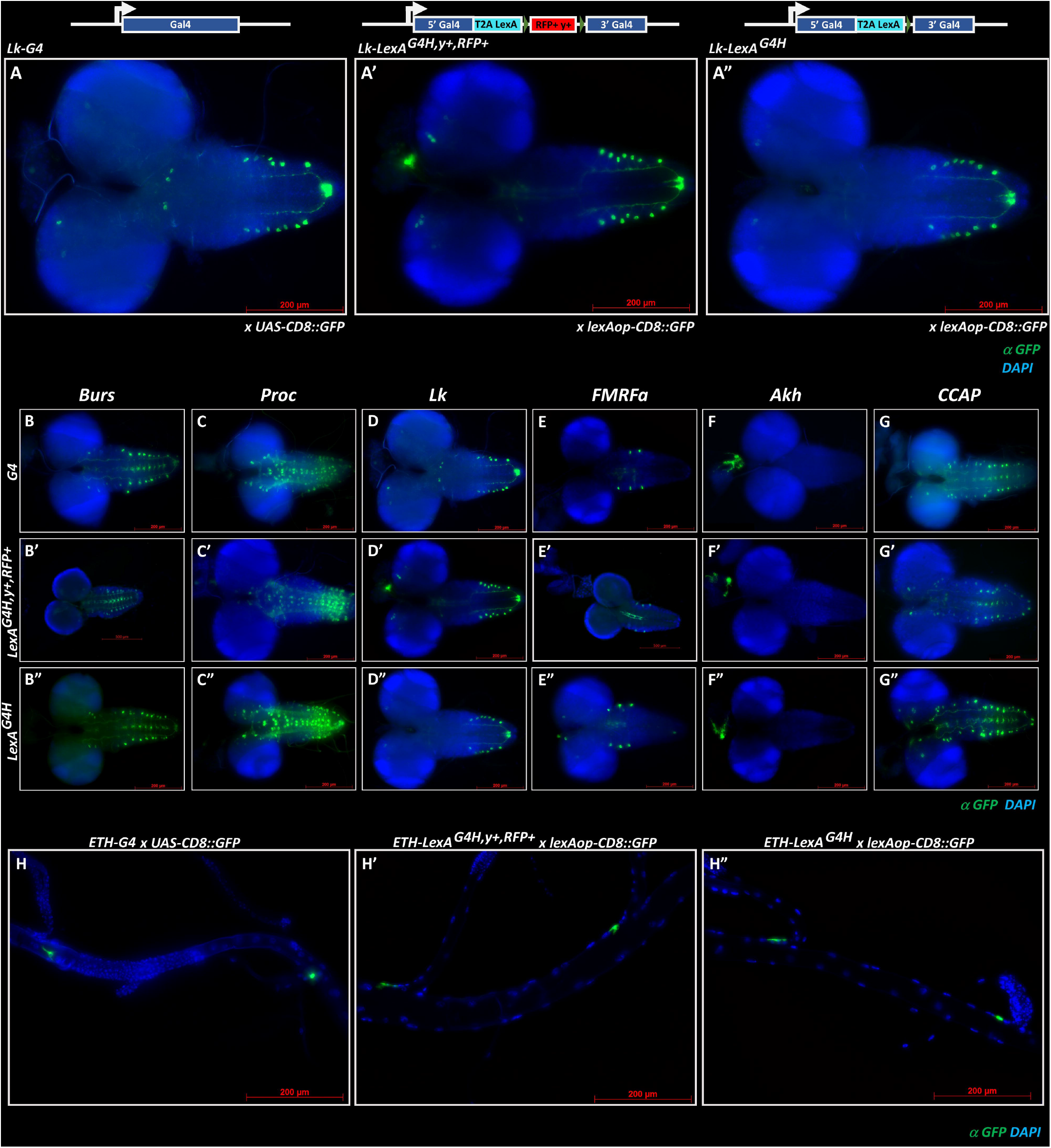
Expression of GFP reporter by parental *Gal4*, the *LexA*^*G4H,y+,RFP+*^ intermediate, and after removal of the *y+,RFP+* selection cassette (*LexA*^*G4H*^) in L3 larval brain and gut. **(A-A”)** Stages of *Lk-Gal4* conversion analyzed by GFP reporter expression (*UAS-CD8*::*GFP* for *Lk-Gal4, LexAop-CD8*::*GFP* for *Lk-LexA*^*G4H,y+,RFP+*^ and *Lk-LexA*^*G4H*^ derivatives). Structure of genomic constructs (see **Figure 1**) are indicated above. The specific driver expression of the *Lk* promoter fragment is present in parental *Gal4*, intermediate *Lk-LexA*^*G4H,y+,RFP+*^ and fully converted *Lk-LexA*^*G4H*^ after the RFP^+^, y^+^ cassette has been removed by hs-Cre. Blue: DAPI. Green: Anti-GFP. **(B-G”)** Conversion of 6 different promoter-*Gal4* constructs analyzed by GFP reporter expression (*UAS*-*CD8*::*GFP* for *Gal4, LexAop-CD8*::*GFP* for *LexA*^*G4H,y+,RFP+*^ and *LexA*^*G4H*^ derivatives). (B-B”) *Burs-Gal4*, (C-C’) *Proc-Gal4*, (D-D”) *Lk-Gal4* as shown in A-A”, (E-E”) *FMRFa-Gal4*, (F-F”) *Akh-Gal4*, (G-G”) *CCAP-Gal4*. The stage of conversion is indicated on the left and as indicated in (A-A’). In these lines, the GFP expression pattern remains unchanged from *Gal4* via *LexA*^*G4H,y+,RFP+*^ to *LexA*^*G4H*^. Blue: DAPI. Green: Anti-GFP. **(H-H”)** Conversion of the Inka-cell specific *ETH-Gal4* line to *ETH-LexA*^*G4H*^. L3 dorsal trunk trachea shows GFP positive cells displaying Inka-cell morphology located near branchpoints of dorsal trunks. (H) *ETH-Gal4*, (H’) *ETH-LexA*^*G4H,y+,RFP*^, and (H”) *ETH-LexA*^*G4H*^. Genotypes indicated on top. Blue: DAPI. Green: Anti-GFP.

Seven of the eleven successfully HACKed *Gal4* lines show equivalency of expression independent of the *y*^+^*RFP*^+^ cassette (**Figure 3**). Parental as well as HACKed derivatives (*LexA*^*G4H,y+,RFP+*^ and *LexA*^*G4H*^) of Burs-*Gal4, Proc-Gal4, Lk-Gal4, FMRFa-Gal4, Akh-Gal4, CCAP-Gal4* and *ETH-Gal4* displayed similar-to-identical reporter gene expression (**Figure 3**) in L3 larval CNS (**Figure 3A-G”**) or tracheae of the dorsal trunk (**Figure 3 H-H”**). In conclusion, these *Gal4* and their HACKed *LexA* line derivatives demonstrated expression equivalency.

### Non-equivalency of parental *Gal4* and HACKed *LexA* expression in strain subsets

Analysis of prior HACKed *Gal4* lines (Rankin et al., 2023) and this study (**Figure 3**) report indistinguishable expression domains of parental *Gal4* and the HACKed derivative *LexA* (“HACKed expression domain equivalency”). Here, we observed two exceptions to this general rule (**Figure 4**):

**Figure 4:**
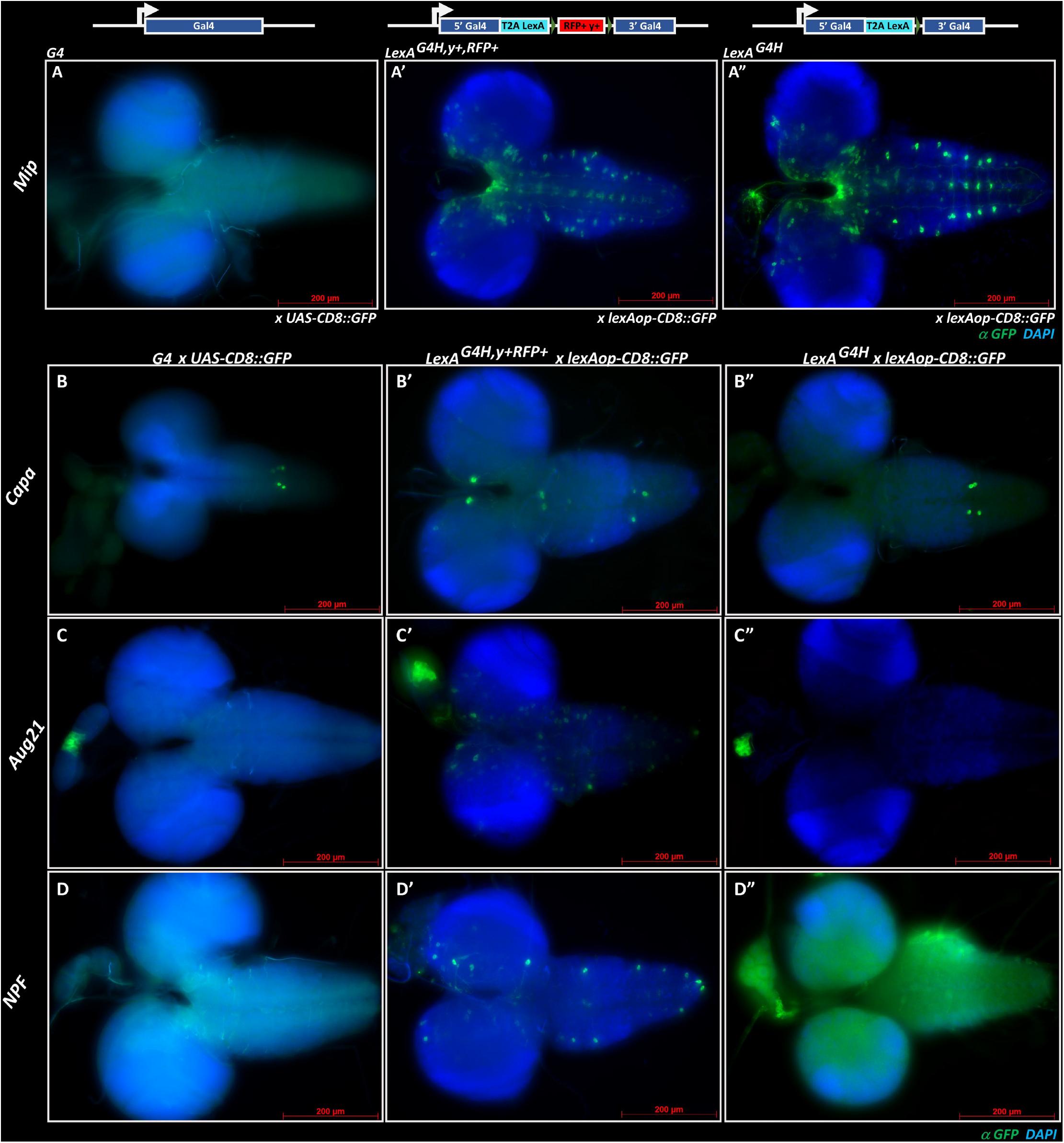
*Gal4* lines with alteration of GFP expression during the *Gal4 -LexA* conversion. **(A-A”)** Converted *Mip-LexA*^*G4H,y+,RFP+*^ and *Mip-LexA*^*G4H*^ show GFP expression derived from a silent parental *Mip-Gal4* promoter construct (absent GFP expression in *Mip-Gal4; UAS-CD8-GFP*). The genomic constructs) are shown on top. Blue: DAPI. Green: Anti-GFP. **(B-C”)** *Gal4 lines showing additional expression domains as intermediate LexA*^*G4H,y+,RFP+*^, *but not as fully converted y*^*-*^,*RFP*^*-*^ *LexA*^*G4H*^. *Capa-LexA*^*G4H,y+,RFP+*^ *(B’) and Aug21-LexA*^*G4H,y+,RFP+*^ *(C’) L3 larval brains display additional GFP positive neurons in CNS and VNC* in comparison to their parental *Gal4* (B,C) and fully converted *LexA*^*G4H*^ (B”,C”). *Genotypes indicated on top. Driver line promoter indicated on left. Blue: DAPI. Green: Anti-GFP*. **(D-D”)** The silent, GFP negative *NPF-Gal4* line (D) shows GFP positive neuronal expression as intermediate *NPF-LexA*^*G4H,y+,RFP+*^ (D’), and returns to silent, GFP negative state as fully converted *y*^*-*^,*RFP*^*-*^ *NPF-LexA*^*G4H*^ (D”, displays green autofluorescence due to long exposure time). Genotypes indicated on top. Driver line promoter indicated on left. Blue: DAPI. Green: Anti-GFP.

1. “Lazarus-Line” (**Figure 4A-A”**). Careful expression analysis of *MIP-Gal4* (BDSC 51984: **Methods**) crossed to *UAS-CD8::GFP* did not reveal detectable reporter line expression during the L3 stage. In contrast, the HACKed MIP-LexA^G4H,y+,RFP+^ and *MIP-LexA*^*G4H*^ derived lines display robust and reproducible expression in CNS and VNC consistent with neuropeptide-expressing neurons (Nässel and Zandawala, 2020; see Discussion).
2. “Marker-cassette dependent ectopic expression” (**Figure 4 B-D”**). Comparison of the *LexA*^*G4H,y+,RFP+*^ versus *LexA*^*G4H*^ derivatives of *Capa-Gal4, Aug-Gal4* and *NPF-Gal4* (**Figure 4 B’, C’ D’, Methods**) revealed additional *GFP*^+^ neurons in L3 CNS and VNC of *LexA*^*G4H,y+,RFP+*^ progeny that might be attributable to the presence of *y*^+^*RFP*^+^ cassette. Removal of the *y*^+^*RFP*^+^ cassette produces an expression domain of *LexA*^*G4H*^ that was indistinguishable from the parental *Gal4* strains. Of note, the transcriptionally inactive *NPF-Gal4* (BDSC 25681) that shows ectopic GFP expression as *NPF-LexA*^*G4H,y+,RFP+*^ is returned to transcriptional inactivity as its *NPF-LexA* ^*G4H*^ descendant. We conclude that the *y*^+^*RFP*^+^ selection cassette might have input into the expression domain of the hosting *Gal4* line. We conclude that careful examination of *LexA* expression in HACKed lines is necessary, especially if the *y*^+^*RFP*^+^ selection cassette remains inserted (**Figure 1**).

### Integrating *Gal4*-to-*LexA* HACKing into summer secondary school curricula

To demonstrate the feasibility of a summer science class (**Figure 5**) that deployed CRISPR/Cas9 mediated gene conversion described here, we executed the program in a seven-week long class at Harvard DCE in 2023 with 12 students, including 10 high school participants. The class consisted of two modules (M): M1 involved the genetics of *Gal4*-to-*LexA* gene conversion that implemented the crossing program to *F*_3_ (**Figure 1B**), M2 contained the analysis of expression domain equivalency between the parental *Gal4* lines, and their *LexA* descendants (**Figure 3, 4**).

**Figure 5:**
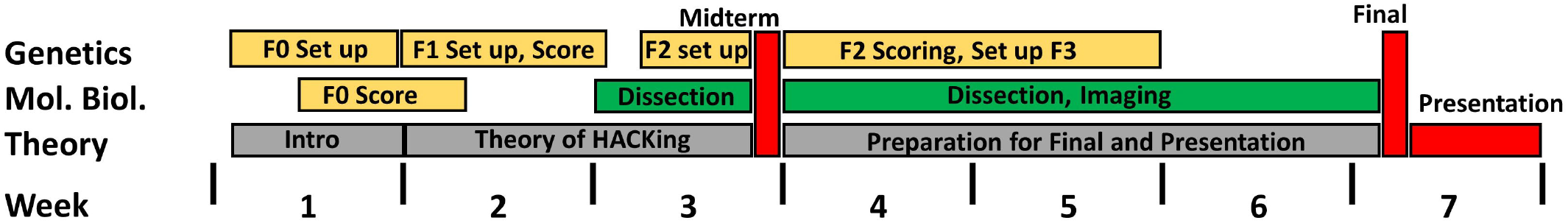
Orchestration of Genetics, Molecular Biology (Mol. Biol.) and Teaching of CRISPR Theory within a seven-week class. Teaching of CRISPR theory took place every day for approximately 60 mins (indicated in gray) for three weeks, and was replaced afterwards by lectures preparing students for the final exam and their presentation (shown in red). The *F*_*3*_ genetics of *Gal4* HACKing (exhibited in yellow, see also **Figure 1**) dominated the hands-on work of the course up to week 3. *F*_*0*_ and *F*_*1*_ were prepared by an instructor in advance to accommodate to the available time. Training in L3 larval microdissection and imaging of reporter line crosses to *LexAop-CD8*::*GFP* and *UAS-CD8*::*GFP* (displayed in green) was emphasized from week 4 onward.

The four-generation mating scheme within a seven-week schedule was achieved by instructor-driven *F*_0_ and *F*_1_ (**Figure 1B**) intercrosses, at four and two weeks before the course commencement, respectively. This enabled students to begin screening *F*_0_ progeny on the first day of class, and *F*_1_ progeny within the first week. *y*^+^*RFP*^+^ cassette removal was undertaken separately after the class had concluded. This index class established: (1) the generation of balanced, ‘genetically stable’ *LexA*^*G4H,y+,RFP+*^ lines in a *y*^1^,*w*^1118^ background, (2) *LexA*^*G4H,y+,RFP+*^-dependent CNS and VNC expression of the *LexAop-CD8::GFP* reporter in parallel to studies establishing *UAS-CD8::GFP* reporter expression of the parental *Gal4* line, (3) examination of the expression equivalency of parental *Gal4* and *HACKy*.*V2*-derived *LexA* lines, and (4) distribution of the generated *Drosophila* lines to the BDSC. In summary, the class established the feasibility of a CRISPR/Cas9-based curriculum for high school or college student implementation and delivery of novel *LexA* lines ready for use by the science community.

## Discussion

To address a global desire for *LexA* lines with cell type-specific expression patterns, we and others conducted enhancer trap screens or a directed cloning approach (Pfeiffer et al 2010; Kockel et al 2016, Kockel et al 2019, Kim et al 2023, Wendler et al 2022). However, the enhancer trapping approach requires extensive molecular characterization, including insertion site and expression mapping. An alternative approach is represented by CRISPR/Cas9 ‘HACKing’, generating *LexA* lines that maintain the previously-characterized insertion site and expression domain of their parental *Gal4* (Lin and Potter 2016; Chang et al 2022, Rankin et al., 2023). We applied the concept of CRISPR/Cas9 mediated *Gal4*-to-*LexA* conversion (Rankin et al., 2023) to 12 neuropeptide-specific *Gal4* lines, with the successful conversion of 11 *Gal4 lines*. The single unsuccessful conversion resulted from mis-mapping of the parental line *AstA-Gal4* (with *Gal4* on chromosome III) to chromosome II (BDSC 51979). Remarkably, none of the >40 individual *F*_1_ crosses of *y*^1^,*w*^1118^ *vas-Cas9; CyOHACKy/+; AstA-Gal4/+* males to *y*^1^*w*^1118^ females yielded a converted male. The chromosomal preference of the *CyO*^*HACKy*.*V2,y+,RFP+*^ donor for converting chromosome II targets with much higher frequency than chromosome III targets is well documented (Rankin et al., 2023; Lin and Potter, 2016), and re-confirmed here. All other *Gal4* drivers were located on chromosome II and readily converted within the crossing scheme. In general, the parental *GAL4* and its conversion derivative *LexA*^*G4H*^ displayed similar-to identical tissue expression patterns, indicating molecular conversion events of high fidelity.

The presence of the selectable *yellow*^+^ cassette within the *HACKy*.*V2,y*^*+*^,*RFP*^*+*^ donor sequence enabled rapid selection of *Gal4*-to-*LexA* conversion events, that could be confirmed by the presence of the *RFP*^+^ marker. The requirement for only a simple stereoscope setup to select for Gal4-to-LexA conversion events by the presence of the *yellow*^+^ coat color marker in male *F*_1_ offspring broadens the feasibility of a HACKing-based curriculum in any classroom setting, and brings significant time-savings for the academic researcher. Thus far, all phenotypically observed *yellow*^+^ *F*_1_ male offspring also displayed expression of RFP in the ocelli, eye, or brain; thus, fluorescent microscopy-based *RFP*^+^ selection could be used to confirm conversion, but is not essential.

The HACKy.V2,*y*^*+*^,*3xP3-RFP*^*+*^ donor contains a synthetic P3 promoter element based on *pax6* upstream regulatory sequence (Sheng et al 1997). Upon *Cas9*-mediated integration into the *Gal4* target, the 3xP3 promoter element reliably drives RFP expression in *Drosophila* ocelli, eye, or brain, and was used as the only selectable marker in a prior version of the HACK donor (Chang et al, 2022). As in prior work (Rankin et al., 2023), we observed that once integrated into the *Gal4* ORF, we observed in some instances a transcriptional effect of the presence of the *3xP3 pax6* promoter on the expression domain of the HACKed *Gal4* line. This was evident in four out of eleven HACKed neuropeptide-*Gal4* lines, *Mip*-*LexA*^*G4H,y+,RFP+*^, *Capa-LexA*^*G4H,y+,RFP+*^, *Aug21-LexA*^*G4H,y+,RFP+*^, and *NPF-LexA*^*G4H,y+,RFP+*^ (**Figure 4**). Cre-mediated deletion of *the y*^*+*^,*RFP*^*+*^ selection cassette eliminated ectopic expression (**Figure 4**). Hence, we strongly advise *y*^*+*^*RFP*^*+*^ selection cassette removal after HACKing.

In one case, we unexpectedly identified a *Gal4* line *(Mip-Gal4*, BDSC 51984) without evident Gal4 activity, from which we HACKed *LexA*^*G4H,y+,RFP+*^ and *LexA*^*G4H*^ derivatives with a robust and reproducible expression pattern in the CNS and VNC of the L3 larval brain, conforming to a known pattern of neuropeptide expression (Nässel, and Zandawala, 2020, **Figure 4A-A”**). This “Lazarus-effect” of revived expression from a silent parental line to transcriptionally active HACKed derivatives might be caused by mutation of the parental *Gal4* ORF that nevertheless allows for homology-assisted repair. Future sequencing efforts could assess this possibility.

The foundation of our science outreach to high schools, an effort called Stan-X (www.Stan-X.org) with 21 participating schools as of 2024, is a modernization of biology curricula, with a focus on teacher training to enable classroom-based unscripted research by student-scientists. To that end, we have developed and implemented the seven-week CRISPR-based genome editing course in *Drosophila* described here. It combines modern science with the generation of novel *LexA* expression tools that are distributed to the scientific community. Our course uses *Drosophila*, a safe, fast and economic genetic model organism that supplements basic theoretical training in mendelian and non-mendelian genetics widespread in most secondary school and university biology curricula. Our programs enable student-scientists to execute experiments and to present data to peers and the community of science, including through oral presentations, and manuscripts or publications (Kockel et al 2016, 2019; Chang et al 2022; Kim et al 2023; Rankin et al 2023). This short course has been ideal for adaptation by Stan-X partner schools that have recently adopted it for instruction in the regular academic year. Reconstituting genuine, experience-based science practice in secondary school settings should be advanced by findings, tools and methods detailed here.

## Acknowledgements

We thank the Bloomington Drosophila Stock Center (NIH P40OD018537) for fly stocks and FlyBase (NHGRI P41HG000739) for updated information. We thank past and current members of the Kim lab for helpful discussions. We are grateful to Philip Weissman (Micro-Optics Precision Instruments, NY) and Ken Fry (Genesee Scientific, CA) for generous support of equipment procurement for this project. We thank the Associate Dean Karen Flood, and Diana H Belanger and Clark Magnan for their efforts to support this course.

